# Variabilities and similarities of adult stem cells derived intestinal organoids originating from different intestinal segments in pig

**DOI:** 10.1101/2022.12.14.520382

**Authors:** Soumya K. Kar, Marinus F.W. te Pas, Roxann Rikkers, Ole Madsen, Nico Taverne, Esther D. Ellen, Jerry M. Wells, Dirkjan Schokker

## Abstract

Organoids are *in vitro* model systems generated from tissues. Organoids express specific physiological functions associated with their original tissue location and they express tissue-segment-specific genes. The aim of this study was to culture pig organoids from different areas of intestinal segments: duodenum, ileum (with or without Peyers Patches (PP)), and colon, to investigate the resemblance with the *in vivo* tissues and variability of multiple adjacent sampling sites based on histology and transcriptome profiles. The transcriptome profiles of the *in vivo* tissues and the derived organoids showed high resemblance for all intestinal segments. For the transcriptomic cluster analysis it was shown that it is important to use tissue important genes to shown the resemble between tissue and their derived organoids. The transcriptome profiles clearly separated the intestinal segments, and samples of the same segment from adjacent tissue locations showed high transcriptome profile similarity. Ileum samples with and without PP were also separated. Pathway analysis of differentially expressed genes from PP compared with non-PP suggested the importance of several aspects of cell cycle progression regulation, including DNA metabolism, chromatin organization, regulation of mitotic stage progression, and regulation of inflammation. Based primarily on the transcriptomics results, we conclude that organoids reflect the sampled intestinal segment and that organoids derived from adjacent sampling sites in an intestinal tissue segment showed low variability. The results from the ileum indicate that organoids have potential to study intestinal innate immune processes.

## 2. INTRODUCTION

The intestine is crucial for digestion and uptake of nutrients, but also provides a barrier between the host and lumen content, which is colonized by commensal microorganisms and sometimes pathogens, including viruses. *In vitro* methods for studying gut functionality can facilitate development of nutritional, prebiotic or probiotic strategies to further improve intestinal and overall health (Wells et al., 2017; Nyachoti and Lee, 2020; Lauridsen, 2020; Yang and Liao, 2019; Pluske et al., 2018). In the last decade, researchers are increasingly adopting adult stem cell (ASC)-derived intestinal organoids from pigs and other animals, as an *in vitro* research tool, to get further insights into gut health and functioning (Kar et al., 2021). The intestinal organoid culture provides many of the heterotypic cell lineages found in the tissue/organ of origin. In addition, the intestinal organoids are both genetically and phenotypically stable during prolonged periods of cell culture and are amenable to standard experimental manipulations. Organoids are therefore promising *in vitro* model systems that recapitulate biological processes and interactions more accurately than non-organoid cell cultures (Van der Hee et al., 2018; Van der Hee, et al., 2020; Kar et al., 2021). However, there are some knowledge gaps associated with the ASC-derived intestinal organoid culture that need to be addressed to ensure reliability and reproducibility of results obtained from such *in vitro* model. As organoids are generated from intestinal ASCs that perform specific functions related to their original segment in the gut, they express segment-specific genes (Middendorp et al., 2014; Kolawole and Wobus, 2020). Nevertheless, there is potential for variability and reproducibility among samples from nearby locations from the same tissue that may be important for their application as *in vitro* models. Low variability in the transcriptome of intestinal organoids obtained from independent jejunum samples from the same animal has been reported (Van der Hee et al., 2020), but further studies are needed for other segments, particularly the ileum. Ileum is lined with Peyers patches (PP) containing agglomerates of numerous lymphoid follicles. In pigs, the lymphoid tissue associated with the ileum is located on one side of the intestine, and this cytokine-rich environment may have imprinted epigenetic changes in ASCs that result in segment-specific variability in the transcriptome and phenotype of the organoid (Machado et al., 2005; Cao et al., 2015). Therefore, organoids derived from parts of the ileum with and without PP need to be compared. The aim of this study was to explore segment-specific identity and variability in different pig intestinal segments from tissue and derived organoids, and to investigate organoid resemblance to the *in vivo* tissue from which they were derived, as well as variability in the organoid cultured from the adjacent sampling site within the intestinal segment of the corresponding tissue, based on histology and transcriptome profiles. We cultured replicates of ASC-derived organoids from tissue samples from the duodenum, ileum (with or without PP), and colon. To investigate the variability of porcine organoids, we compared the transcriptome profiles of organoids within and among various intestinal tissue segments. To investigate the resemblance with the *in vivo* situation, we compared the transcriptome profiles cultured organoids with the corresponding intestinal tissue segment. Because of its involvement in immune processes, we paid special attention to ileum segments with and without PPs.

## 3. METHODS

Detailed information of the animal material used, isolation procedures, and organoid characterization with histology and transcriptomics can be found in the supplementary information. Briefly, three adjacent tissue samples from the duodenum, ileum, and colon of a 19-day male piglet were collected. In the ileum samples both PP and non-PP-associated tissues were collected. Epithelial crypt cells were plated in Matrigel (Corning, Corning NY, USA) coated culture dishes to grow multi cell type 3D organoids from stem cells.

Histomorphometric characterization was carried out for both native and cultured organoids. Organoid cultures were frozen after ten days. The whole-genome transcriptome landscape was measured in both intestinal organoids and intestinal segment using RNAseq (Novogene, Milton Road, Cambridge, UK). The transcriptomics analysis pipeline is described in the Supplementary Figure S1. Results of transcriptome profiles intestinal organoid cultures were compared to native intestine tissues. Detailed information of the animal material used, isolation procedures, and organoid characterization with histology and transcriptomics can be found in the supplementary information. The analysis pipeline is described and visualized in Supplementary Figure S1.

## 4. RESULTS

Histomorphometric analysis of the ileum tissue sections revealed that the lymphoid tissue was located only on the outside of the intestine (Supplementary Figure S2). In areas containing PP, immune cells were close to the follicular-associated epithelium and a higher number of immune cells were present in the lamina propria compared to non-PP mucosa.

Supplementary Table S1 shows the analyses and statistics of the transcriptome data. Transcriptome profiling, clustering (Figure 1A), and the correlation matrix (Figure 1B) separated the intestinal segments and derived organoids, which showed high similarity between segment tissue replicates and their derived organoid replicates (Supplementary Figure S3). Figure 1C shows a heatmap of important intestinal genes (details can be found in Supplementary Table S2) measured in the four intestinal segments and their derived organoids. These results (Figure 1A, B, C and Figure S3) seem to indicate that the tissues and the organoids have distinct expression patterns where the organoids tend to cluster together, instead of clustering with the tissue of origin.

**Figure 1.**
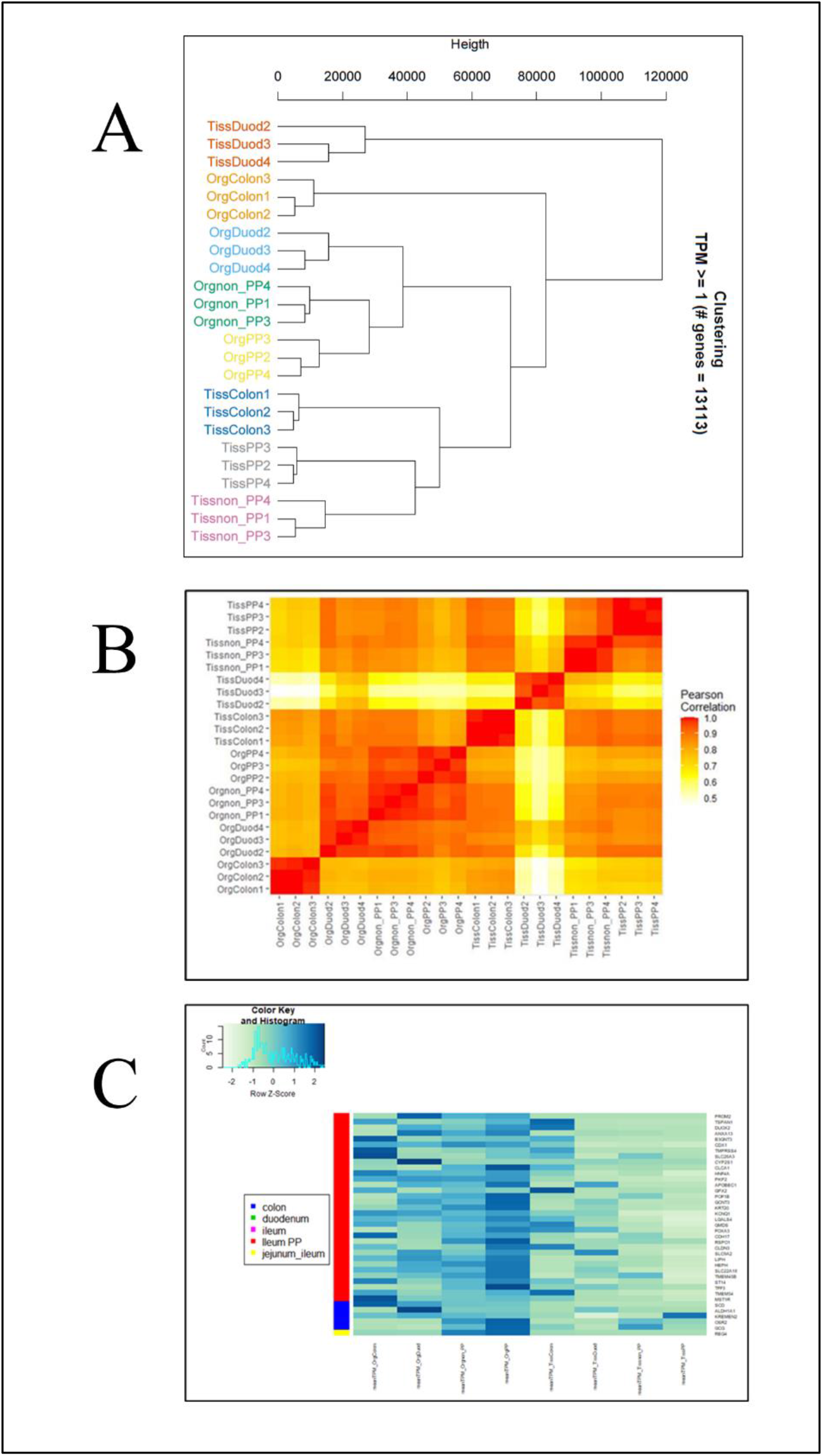
Clustering (A), correlation matrix (B), and Heatmap (C) based on the transcriptomics profiles of organoids and intestinal segments. Figure C was prepared with genes showing differential expression between the intestinal tissues and derived organoids.

Cluster analysis with the transcriptome profile of the important intestinal genes (Supplementary Table S2) clearly showed that the organoids of the colon and duodenum resemble the corresponding tissue from which they are derived (Figure 2A). The PCA analysis (Figure 2B) showed that principal component 1supported the similarity of organoids and tissue. Principal component 2 showed that the PP and non-PP tissues and organoids cluster together. Together this showed that the organoids resembled the tissues they were derived from when using important intestinal genes. Furthermore, genes expressed higher in tissues at a specific intestinal segment showed a similar higher expression in the derived organoids, suggesting a more direct relation between the tissue and the derived organoids. This relation between tissue and derived organoids was not obvious from the global gene expression. Furthermore, both clustering and PCA plots (Figures S3A and S3B, respectively) suggest that duodenum (PCA first component) and colon organoids (PCA second component) are distinct from the other samples both for tissue and organoids.

**Figure 2.**
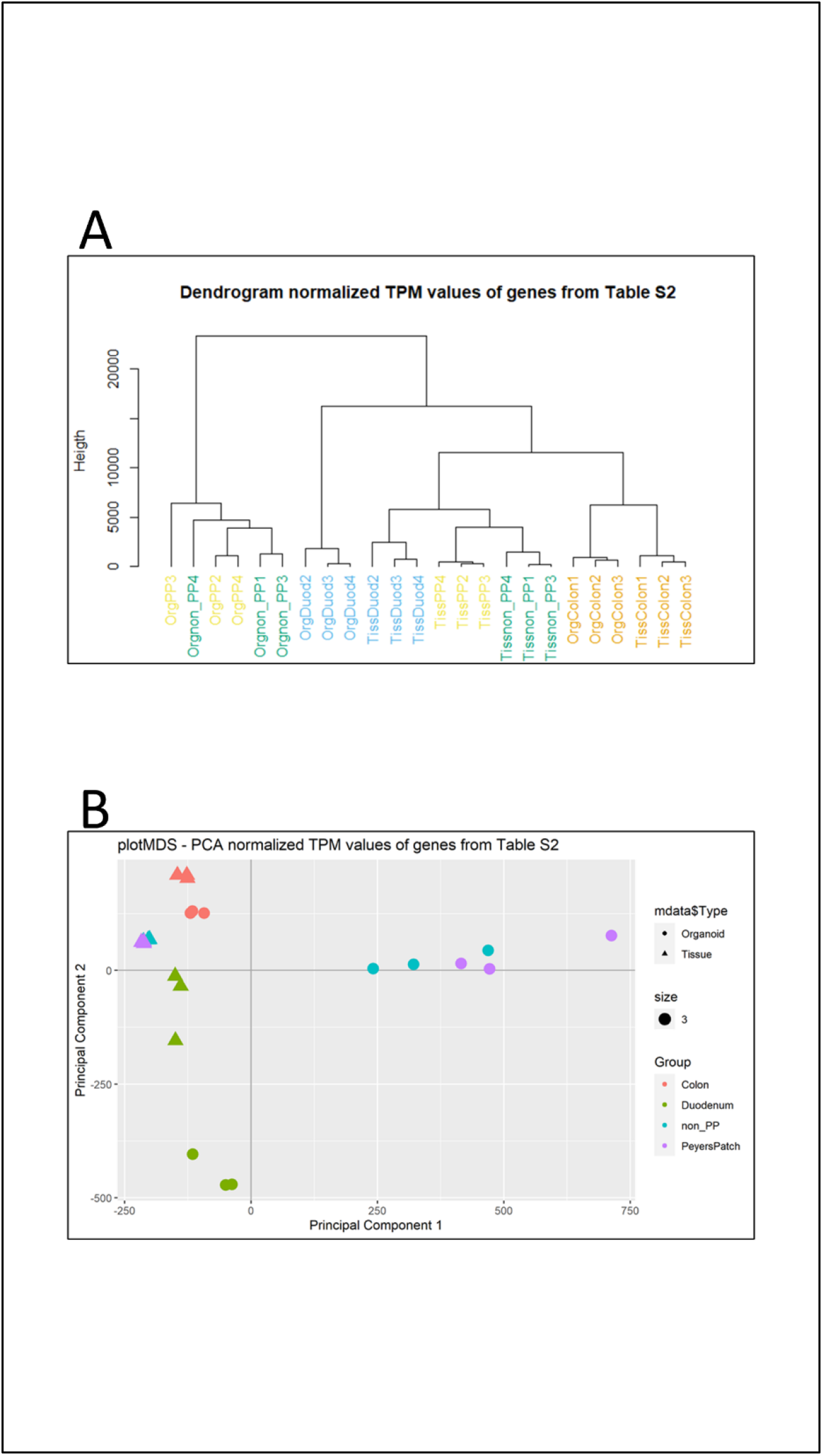
Dendrogram (A) and Principal component analysis (B) of the genes from Table S2 indicating the resemblance of the tissues and their derived organoids. The Figure shows clustering of the tissues and their derived organoids of colon and duodenum with Principal component 1, and clustering of PP and non-PP with Principal component 2.

Organoids and tissue transcriptome profiles share large parts of the transcriptome profile (Supplementary Figure S4). For all intestinal segments, approximately 80% of the genes are expressed in both tissue and organoid segments. For duodenum the correlation tissue-organoid is approximately 0.7, and for colon, ileum-PP and ileum-non-PP >0.8 (Figure 1B). The number of genes not shared between tissue and organoids is always higher in the tissue compared with the organoids, although organoids also showed some unique gene expressions.

For the ileum of *in vivo* tissue (Figure 3A) and derived organoids (Figure 3B) pathway analyses showed the specific contribution of the PP to the transcriptome profiles. Figure 3C shows the pathways related to differentially expressed genes. The enrichment analysis of differentially expressed genes between PP and non-PP, showed that, although the specific pathways vary, the observed biological processes were similar: (1) Both the *in vivo* tissue and organoids showed regulation of the mitotic cell cycle via regulation of the organization of the chromosomes and DNA replication at various stages, and (2) B-cell mediated adaptive immune response and inflammation. One remarkable difference between tissue and organoid pathway profiles was that the regulation of the mitotic cell cycle and the corresponding DNA synthesis and chromatid organization are positively regulated in organoids and negatively regulated in the tissues. Furthermore, especially in organoids we observed pathways related to the organization and biogenesis of cellular components such as organelles, which we did not observe in the tissue. The data in the Supplementary Figure S4 and Table S2 shows that many genes participate in pathways related to similar processes (for a gene list see supplementary Table S2).

**Figure 3.**
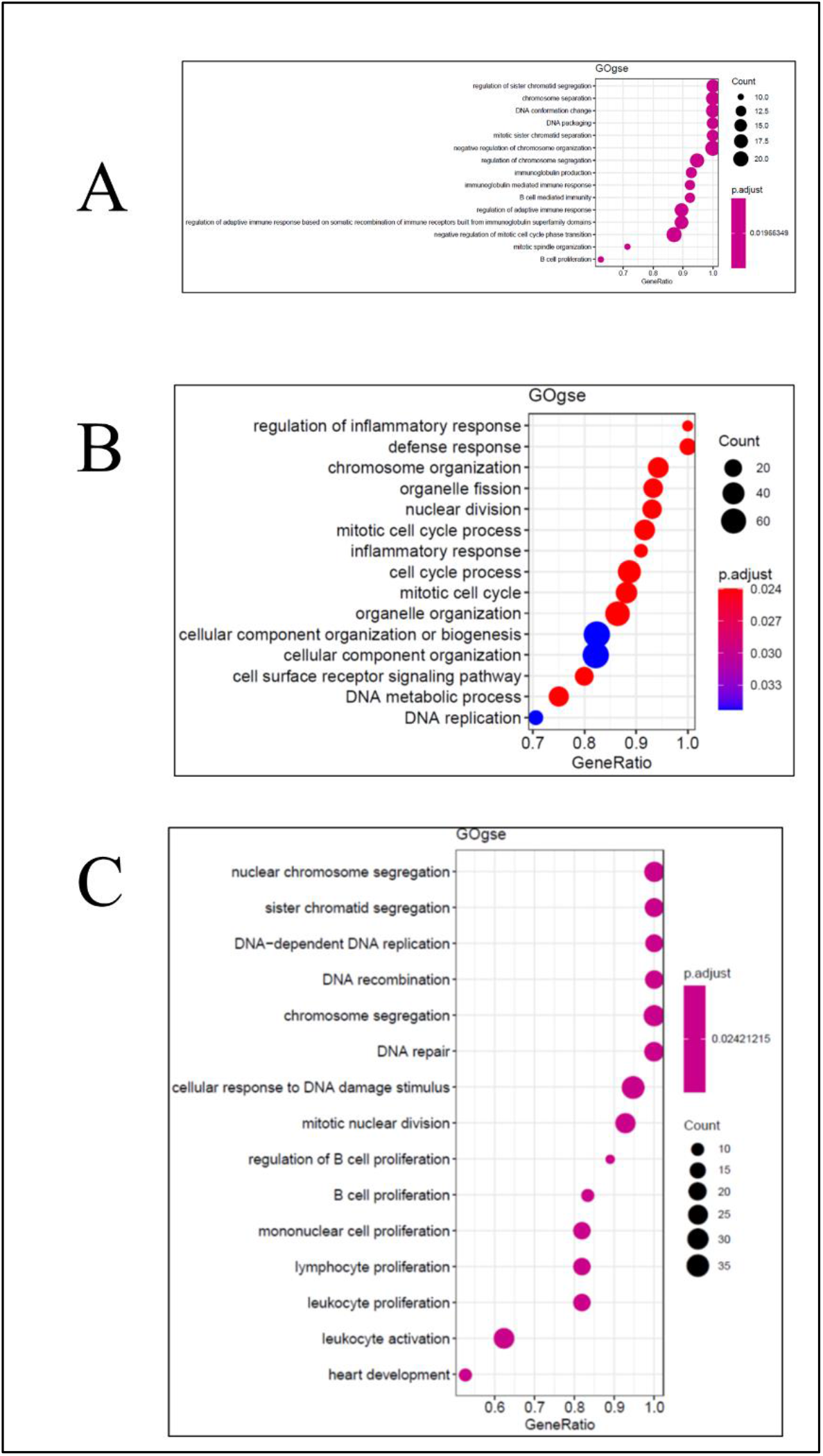
Pathways analysis of the transcriptome profiles of the ileum *in vivo* tissue (A) and organoids (B) comparing samples with and without Peyers Patches (PP). Supplementary Figure S4 shows information about the number of overlapping and non-overlapping genes. (C) Pathways related to differentially expressed genes.

## 5. DISCUSSION

Our results showed that transcriptome profiles of the organoids showed good resemblance to the *in vivo* tissues in the pig intestine. For duodenum the correlation tissue-organoid is approximately 0.7, and for colon, ileum-PP and ileum-non-PP >0.8. However, when comparing the global expression of genes, organoids from different intestinal segments group more together compared to their respective tissue of origin. This grouping of organoids could indicate a specific difference between the *in vivo* and *in vitro* situation in the overall expression of genes, relating to the continuous growth and development that organoids go through during organoid development from intestinal stem cells (Bolhaqueiro et al., 2018; Hausmann et al., 2020). This may relate to the continuous growth of the organoids compared to only tissue renewal of the ileum tissue supported by the functional enrichment of e.g. chromatin organization and DNA packaging pathways, which underline the importance of this finding. The growth and development of organoids is further underlined by the cellular component organization pathway.

Organoids from different intestinal segments were separated and clustered with the corresponding tissue based on important intestinal genes. This shows that intestinal organoids are a good *in vitro* model that resembles the tissue from which they were derived, especially in terms of gene expression. This suggests that the specific transcriptome profile described better the tissue functionality in organoids, and that global transcriptome cluster comparisons alone are not well suited to determine specific cell types found in organoids (Lu et al., 2021). Therefore, further analyses are needed to explore the underlying biological mechanisms behind the general clustering of organoids and on how to establish gene expression profiles that can determine cell composition of organoids. Such an approach could be single cell transcriptomics (Tanaka et al., 2020).

In this study, we found that the transcriptome profile of organoids obtained from intestinal segments represented the tissue well, i.e. similarities in expressed genes among tissue locations and derived organoids is over 80%. We concluded that low precision tissue sampling within an intestinal site yields organoids that are a good representation of the tissue in terms of transcriptome profile. Alternatively, genes specific/indicative for a segment could provide information about the similarities of organoids to the tissue from which they originate. Not all genes were shared between tissue and organoids. The number of genes not shared between tissue and organoids was always higher in the tissue. This indicates the complexity of the tissue, which is higher compared to the complexity of the organoids (Achberger et al., 2019). The tissue contains smooth muscle tissue (blood vessels), immune cells, and lymph vessels, which are lacking in the cultured organoids. Alternatively, culture conditions may induce small differences as compared to the *in vivo* situation, such as the specific response to the culture media and matrix in which the organoids were cultured.

Furthermore, we provided conclusive evidence that the transcriptomics profile of organoids obtained from tissue samples taken from adjacent sites of a given intestinal segment has a high correlation coefficient of 0.97 - 1.00. This indicates that organoids obtained from adjacent sites of a given intestinal segment are similar in terms of their transcriptomic profiles. Our results suggest that researchers can collect tissue samples from adjacent sites within an intestinal segment to obtain intestinal organoids that resemble the segment from which they were obtained. Especially on larger animals it may be more difficult to gather repeated samples on an exact location using the biopsy technique. Nevertheless, our results suggest that the organoids technique can be used on pigs represented by a single biopsy, and this conclusion holds for adjacent locations as well.

For the ileum we sampled both locations with and without PPs, based on observational morphological differences (see Figure S2). Our results indicated that locations with and without PPs were also different on the transcriptomic level, which was also observed for the derived organoids. Comparing tissue and organoids showed that the organoids represented the sampling location well, both at PP and non-PP locations. Thus, for the ileum it is important to determine the presence of PPs before culturing. The pathway analysis revealed high similarities between the comparison of non-PP vs. PP in ileum tissue and ileum-derived organoids. However, some specific differences were noted. For example, both expression profiles exhibit cell cycle regulation, but with positive regulation in organoids and negative regulation in tissue. This may relate to the higher need for growth in organoids as compared with intestinal tissue.

The regulation of the inflammatory response pathway in organoids further indicates the similarity to its respective *in vivo* tissue. In the Gene Ontology (Biological Processes) and KEGG pathways database regulation of inflammation indicates a total of 13 pathways. Most important are chemokine synthesis and attraction of leukocytes to the site of inflammation. Some of them can be found in the gene list like LECT2 (Supplementary Table S2). However, this gene list is specific for the intestine while such genes can also exert more general immune functions (Pott and Horneff, 2012). This result points to the high complexity of the organoid organization and the resemblance to the *in vivo* tissue.

Concluding, we showed that transcriptome analysis of tissue-specific genes showed that the organoids resembled the tissues they were derived from. Furthermore, from this study we conclude that it is possible to study intestinal functionality of the duodenum, ileum, and colon segments using a sampling from adjacent location for each of these intestinal segments. It is not necessary to determine the exact location of the sampling site within the intestine segment. For the ileum it is important to determine the presence of PPs before culturing. In this study we used one piglet at one age. It will be necessary to validate the results with more pigs and at different ages.

## Ethics statement

The animal study was reviewed and approved by Animal Care and Use Committee permission-AVD401002015265, 2016.D-0062.027, and 2016.D-0062.040 of Wageningen University and Research, The Netherlands.

## Notes

### Competing Interest Statement

The authors have declared no competing interest.

## References

Achberger, K., Probst, C., Haderspeck, J., Bolz, S., Rogal, J., Chuchuy, J., Nikolova, M., Cora, V., Antkowiak, L., Haq, W., Shen, N., Schenke-Layland, K., Ueffing, M., Liebau, S., Loskill, P. (2019). Merging organoid and organ-on-a-chip technology to generate complex multi-layer tissue models in a human retina-on-a-chip platform. Elife, 8, e46188, https://doi.org/10.7554/eLife.46188.

Bolhaqueiro, A.C., van Jaarsveld, R.H., Ponsioen, B., Overmeer, R.M., Snippert, H.J.,Kops, G.J. (2018). Live imaging of cell division in 3D stem-cell organoid cultures. Meth. Cell Biol. 145, 91–106.

Cao, L., Kuratnik, A., Xu, W., Gibson, J. D., Kolling IV, F., Falcone, E.R., Ammar, M., Van Heyst, M.D., Wright, D.L., Nelson, C.E., Giardina, C. (2015). Development of intestinal organoids as tissue surrogates: cell composition and the epigenetic control of differentiation. Mol. Carcinog. 54, 189–202.

Hausmann, A., Russo, G., Grossmann, J., Zünd, M., Schwank, G., Aebersold, R., Liu, Y., Sellin, M.E., Hardt, W.D. (2020). Germ-free and microbiota-associated mice yield small intestinal epithelial organoids with equivalent and robust transcriptome/proteome expression phenotypes. Cell. Microbiol. 22(6), e13191.

Kar, S.K., Wells, J.M., Ellen, E.D., Te Pas, M.F.W., Madsen, O., Groenen, M A.M., Woelders, H. (2021). Organoids: a promising new in vitro platform in livestock and veterinary research. Vet. Res. 52, 1–17.

Kolawole, A.O., Wobus, C.E. (2020). Gastrointestinal organoid technology advances studies of enteric virus biology. PLoS Pathog. 16(1), e1008212.

Lauridsen, C. (2020). Effects of dietary fatty acids on gut health and function of pigs pre-and post-weaning. J. Anim. Sci. 98(4), skaa086.).

Lu, J., Krepelova, A., Rasa, S.M.M., Annunziata, F., Husak, O., Adam, L., Nunna, S., Neri, F. (2021). Characterization of an in vitro 3D intestinal organoid model by using massive RNAseq-based transcriptome profiling. Sci. Rep. 11, 1–14.

Machado, J.G., Hyland, K.A., Dvorak, C.M., Murtaugh, M.P. (2005). Gene expression profiling of jejunal Peyer’s patches in juvenile and adult pigs. Mamm. Genome, 16, 599–612.

Middendorp, S., Schneeberger, K., Wiegerinck, C.L., Mokry, M., Akkerman, R.D., van Wijngaarden, S., Clevers, H., Nieuwenhuis, E.E. (2014). Adult stem cells in the small intestine are intrinsically programmed with their location-specific function. Stem cells. 32, 1083–1091.

Nyachoti, M., Lee, J. (2020). 13 Opportunities for using low-protein diets for weanling pigs to improve intestinal health. J. Anim. Sci. 98, suppl. 3, 18–19.

Pluske, J.R., Turpin, D.L., Kim, J.C. (2018). Gastrointestinal tract (gut) health in the young pig. Anim. Nutr. 4, 187–196. doi: 10.1016/j.aninu.2017.12.004.

Pott, J., Hornef, M. (2012). Innate immune signalling at the intestinal epithelium in homeostasis and disease. EMBO reports. 13, 684–698.

Tanaka, Y., Cakir, B., Xiang, Y., Sullivan, G.J., Park, I.H. (2020). Synthetic analyses of single-cell transcriptomes from multiple brain organoids and fetal brain. Cell Rep. 30, 1682–1689.

Van der Hee, B., Loonen, L.M.P., Taverne, N., Taverne-Thiele, J.J., Smidt, H., Wells, J.M. (2018). Optimized procedures for generating an enhanced, near physiological 2D culture system from porcine intestinal organoids. Stem Rell Res. 28, 165–171.

Van der Hee, B., Madsen, O., Vervoort, J., Smidt, H., Wells, J.M. (2020). Congruence of transcription programs in adult stem cell-derived jejunum organoids and original tissue during long-term culture. Front. Cell Dev. Biol. 8, 375.

Wells, J.M., Brummer, R.J., Derrien, M., MacDonald, T.T., Troost, F., Cani, P.D., Theodorou, V., Dekker, J., Méheust, A., de Vos, W.M., Mercenier, A., Nauta, A., Garcia-Rodenas, C.L. (2017). Homeostasis of the gut barrier and potential biomarkers. Am. J. Physiol. - Gastrointest. Liver Physiol. 312, G171–G193. doi: 10.1152/ajpgi.00048.

Yang, Z., Liao, S.F. (2019). Physiological Effects of Dietary Amino Acids on Gut Health and Functions of Swine. Front. Vet. Sci. 6:169. doi: 10.3389/fvets.2019.00169.

