## Supplementary information for "Variabilities and similarities of adult stem cells derived intestinal organoids originating from different intestinal segments in pig"

### Variabilities and similarities of adult intestinal stem cells derived organoids related to the tissue site of different intestinal segments in pigs.

Soumya K. Kar[1], Marinus F.W. te Pas[1*], Roxann Rikkers[1], Ole Madsen[2], Nico Taverne[3], Esther D. Ellen[1], Jerry M. Wells[3], Dirkjan Schokker[1]

*1. Wageningen Livestock Research, Wageningen University and Research, Wageningen, The Netherlands; 2. Animal Breeding and Genomics, Wageningen University and Research, Wageningen, The Netherlands; 3. Host-Microbe Interactomics, Wageningen University and Research, Wageningen, The Netherlands*

### Supplementary Materials and Methods

*Animal material*

All animal handling was done according to the Dutch laws on experimental handling. Animal material was derived from an earlier approved animal experiment (Animal Experimental Committee permission AVD401002015265, 2016.D-0062.027, and 2016.D-0062.040). A 19 day old boar was sacrificed. The intestine was removed, and four adjacent tissue samples were collected from the duodenum, ileum, and colon. Peyer's Patches (PP) are easily recognized in the ileum and both PP and non-PP-associated tissue was collected for organoid cultures.

*Crypt isolation from intestinal tissues and organoid culture*

Intestinal samples were scraped to isolate the epithelial crypt cells that were plated in Matrigel (Corning, Corning NY, USA) coated culture dishes using DMEM culture medium supplemented with growth factors as previously described (van der Hee et al., 2018). Stem cells developed into multi cell type 3D organoids. All intestinal organoids were split every 5 days when ~80% of the organoids were larger than 50 µm, and the splitting ratio was between 1:2 and 1:6. Organoids were first disrupted by mechanical shearing by repeated pipetting. If the organoids were not disrupted they were treated with TrypLE (Gibco) for 10 min at 37 $^{o}$C. Cultures were frozen after 10 days.

*Tissues and Organoid characterization with histology and transcriptomics*

Organoid cultures were characterized, compared to *in vivo* intestine tissue, and organoids from the same and different intestine sections, were compared using RNAseq and histology. Intestinal ileum tissue was fixed in 4% PFA and stained with hematoxylin and eosin (HE) or Crossmon staining (1937). Specific emphasis was put to PP recognition in Ileum samples.

Whole-genome transcriptome profiles were generated with RNAseq and the raw data will become available ENA upon acceptance of the paper. Trim Galore version 0.6.6 (Krueger, 2018) with Cutadapt version 1.16 (Martin, 2011) with default settings except for length 35 (default 20) and stringency 6 (default 1), were used to trim low-quality data, remove poly A tails and remove Illumina sequencings adapters. Only paired-end reads where both reads were ≥ 35 bp were used. The trimmed reads were aligned against the pig reference genome and gene annotation (Ensembl Sus scrofa 11.1.103, (Howe et al., 2021) using STAR version 2.7.3a and default settings (Dobin et al., 2013). RSEM version 1.3.0 (Li and Dewey, 2011) with default settings except for strand specific protocol, which was set to 0 to derive all upstream reads from the reverse strand, was used to estimate expected counts and Transcripts per Million (TPM). In the downstream analysis a TPM threshold of ≥ 1 was used for the heatmap profiles and distance plots (R package limma version 3.40.6 (Ritchie et al., 2015)), Principal Component Analysis (PCA), dendrogram clustering (R package stats version 3.6.1 (R Core Team, 2019)) and heatmaps of the gene expression profiles (R package gplots version 3.0.3, (Warnes et al., 2020) in RStudio version 3.6.1 (R Studio Team, 2016).

To determine Differentially Expressed Genes (DEGs) the RSEM expected counts were used with DESeq2 version 1.24.0 (Love et al., 2014; Anders, 2014) and a False Discovery Rate (FDR) < 0.05 according to the Benjamini-Hochberg (BH) correction (Benjamini and Hochberg, 1995). Gene Ontology Biological Processes (GO:BP) and Kyoto Encyclopedia of Genes and Genomes (KEGG) enrichment analyses were determined using ClusterProfiler version 3.12.0 (Yu et al., 2012) and a FDR < 0.05 using the BH correction. For data handling and visualization, respectively R package dplyr version 0.8.5 (Wickham et al., 2020) and ggplot2 version 3.3.1 (Wickham, 2016) were used. For an overview and visualization see Figure S1.

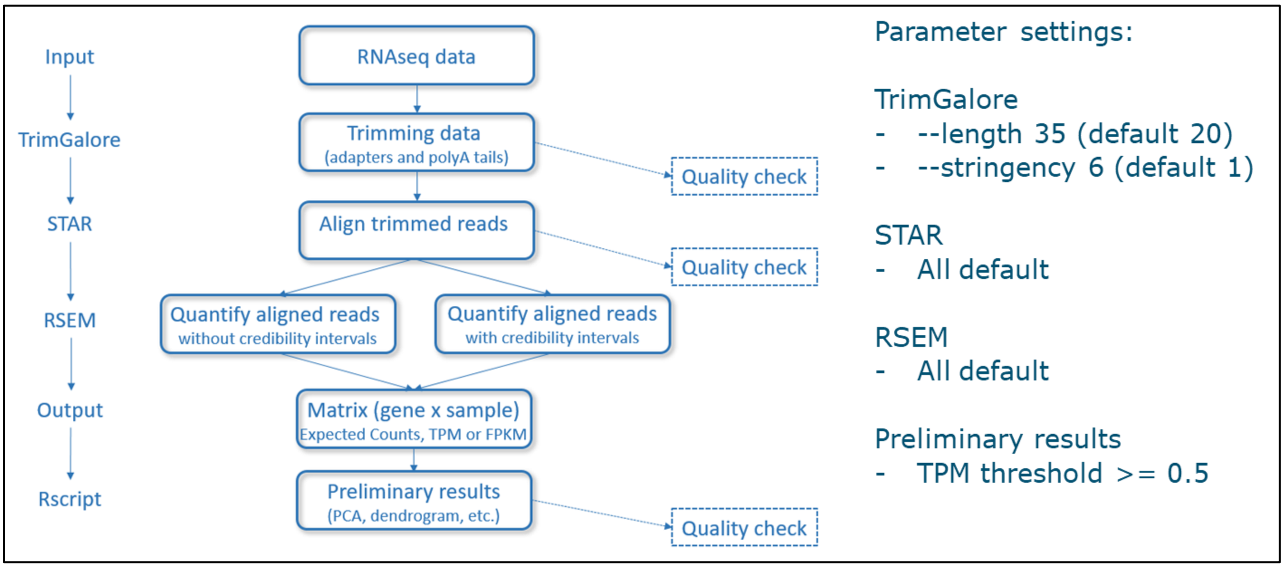

**Supplementary Figure S1**
Visualization of the RNAseq pipeline.

### References

Benjamini, Y., Hochberg, Y. (1995). Controlling the false discovery rate: a practical and powerful approach to multiple testing. J. R. Stat. Soc. series B (Methodological). 57, 289-300.

Crossmon, G. (1937). A modification of Mallory's connective tissue stain with a discussion of the principles involved. Anat. Rec. 69, 33-38.

Dobin, A., Davis, C.A., Schlesinger, F., Drenkow, J., Zaleski, C., Jha, S., Batut, P., Chaisson, M., Gingeras, T.R. (2013). STAR: ultrafast universal RNA-seq aligner. Bioinform. 29, 15-21.

Howe, K.L., Achuthan, P., Allen, J., Allen, J., Alvarez-Jarreta, J., Amode, M.R., Armean, I.M., Azov A.G., Bennett, R., Bhai, J., Billis, K., Boddu, S., Charkhchi, M., Cummins, C., Da Rin Fioretto, L., Davidson, C., Dodiya, K., El Houdaigui, B., Fatima, R., Gall, A., Giron, C.G., Grego, T., Guijarro-Clarke, C., Haggerty, L., Hemrom, A., Hourlier, T., Izuogu, O.G., Juettemann, T., Kaikala, V., Kay, M., Lavidas, I., Le, T., Lemos, D., Martinez, J.G., Marugán, J.C., Maurel, T., McMahon, A.C., Mohanan, S., Moore, B., Muffato, M., Oheh, D. N., Paraschas, D., Parker, A., Parton, A., Prosovetskaia, I., Sakthivel, M.P., Abdul Salam, A.I., Schmitt, B.M., Schuilenburg, H., Sheppard, D., Steed, E., Szpak, M., Szuba, M., Taylor, K., Thormann, A., Threadgold, G., Walts, B.,

Winterbottom, A., Chakiachvili, M., Chaubal, A., De Silva, N., Flint, B., Frankish, A., Hunt, S.E., IIsley, G.R., Langridge, N., Loveland, J.E., Martin, F.J., Mudge, J.M., Morales, J., Perry, E., Ruffier, M., Tate, J., Thybert, D., Trevanion, S.J., Cunningham, F., Yates, A.D., Zerbino, D.R. Flicek, P. (2021). Ensembl 2021. Nucl. Acid. Res. 49, D884–D891.

Krueger, F. (2018). Trim Galore v0.6.6. Retrieved from https://www.bioinformatics.babraham.ac.uk/projects/trim_galore/

Li, B., Dewey, C.N. (2011). RSEM: accurate transcript quantification from RNA-Seq data with or without a reference genome. BMC Bioinform. 12, 323. https://doi.org/10.1186/1471-2105-12-323

Love, M.I., Huber, W., Anders, S. (2014). Moderated estimation of fold change and dispersion for RNA-seq data with DESeq2. Genome Biol. 15(12):550

Martin, M. (2011). Cutadapt removes adapter sequences from high-througput sequencing reads. EMBnet. J. 17, 10-12.

R Core Team (2019). R: A language and environment for statistical computing. R Foundation for Statistical Computing, Vienna, Austria. URL https://www.R-project.org/.

R Studio Team (2016). RStudio: Integrated Development for R. RStudio, Inc., Boston, MA URL http://www.rstudio.com/.

Ritchie, M.E., Phipson, B., Wu, D., Hu, Y., Law, C.W., Shi, W., Smyth, G.K. (2015). Limma powers differential expression analyses for RNA-sequencing and microarray studies. Nucl. Acid. Res. 43(7), e47.

Van der Hee, B., Loonen, L.M.P., Taverne, N., Taverne-Thiele, J.J., Smidt, H., Wells, J.M. (2018). Optimized procedures for generating an enhanced, near physiological 2D culture system from porcine intestinal organoids. Stem Cell Research, 28, 165-171.

Warnes, G.R., Bolker, B., Bonebakker, L., Gentleman, R., Huber, W., Liaw, A., Lumley, T., Maechler, M., Magnusson, A., Moeller, S., Schwartz, M., Venables, B. (2020). gplots: Various R Programming Tools for Plotting Data. R package version 3.0.3. https://CRAN.R-project.org/package=gplots.

Wickham, H. (2016). ggplot2: Elegant Graphics for Data Analysis. Springer-Verlag New York.

Wickham, H., François, R., Henry, L., Müller, K. (2020). dplyr: A Grammar of Data Manipulation. R package version 0.8.5. https://CRAN.R-project.org/package=dplyr.

Yu, G., Wang, L-G., Han, Y., He Q-Y. (2012). ClusterProfiler: an R package for comparing biological themes among gene clusters. OMICS J. Integr. Biol. 16, 284-287

### Supplementary Figures Results

#### Histological analysis

Histomorphometric analysis of the ileum tissue sections revealed asymmetry in the location of the Peyers Patches (PP) and the mucosal-associated lymphoid follicles (Fig S2). The lymphoid tissue was located only on one side and visually apparent as white patches on the outside of the intestine. In areas containing PP immune cells were close to the follicular-associated epithelium and greater number of immune cells were present in the lamina propria compared to the Non-PP ileal mucosa.

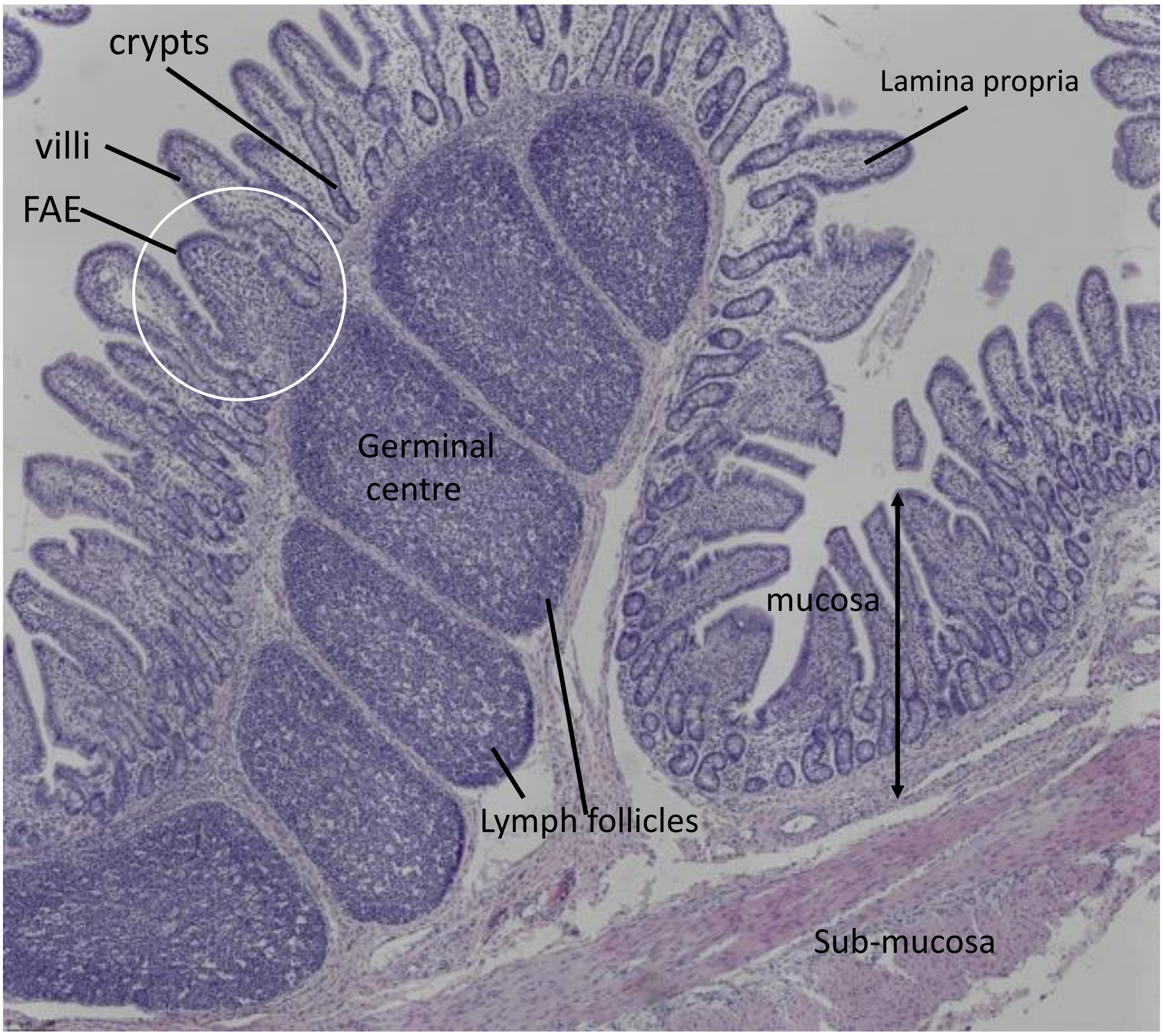

Figure S2.
Ileum from an adult pig. Ileum with Peyers Patches and the epithelial layer above. Above the small PP like domes there is a single cell epithelial layer. Note the lymphoid cells to the basal side of the epithelium. Magnification: 10x.

**RNAseq data**

**A**

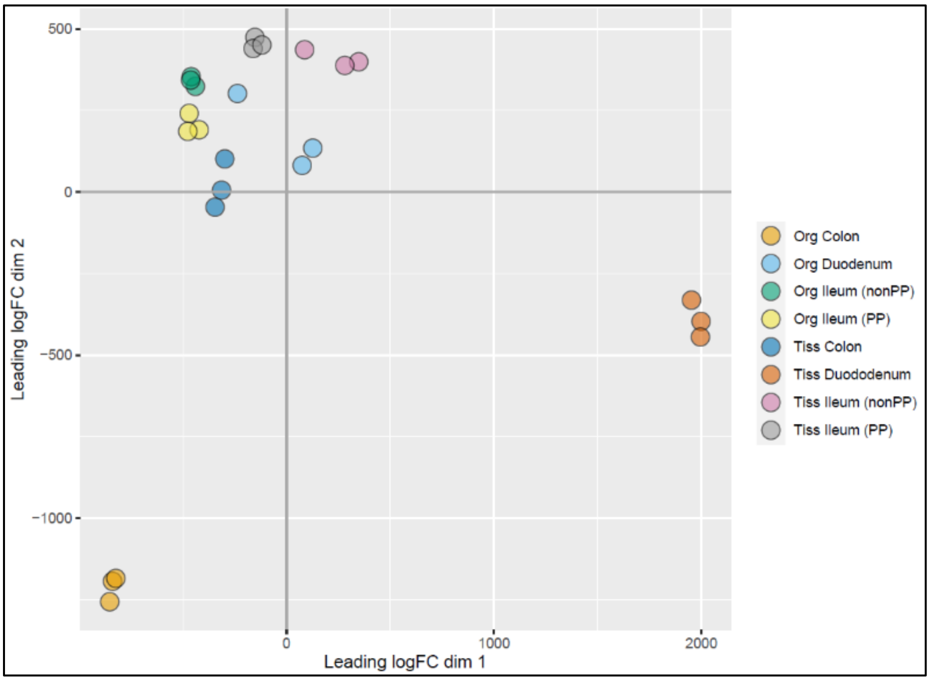
500
0
−500
−1000
0
1000
2000
Leading logFC dim 2
Leading logFC dim 1
Org Colon
Org Duodenum
Org Ileum (nonPP)
Org Ileum (PP)
Tiss Colon
Tiss Dododenum
Tiss Ileum (nonPP)
Tiss Ileum (PP)

**B**

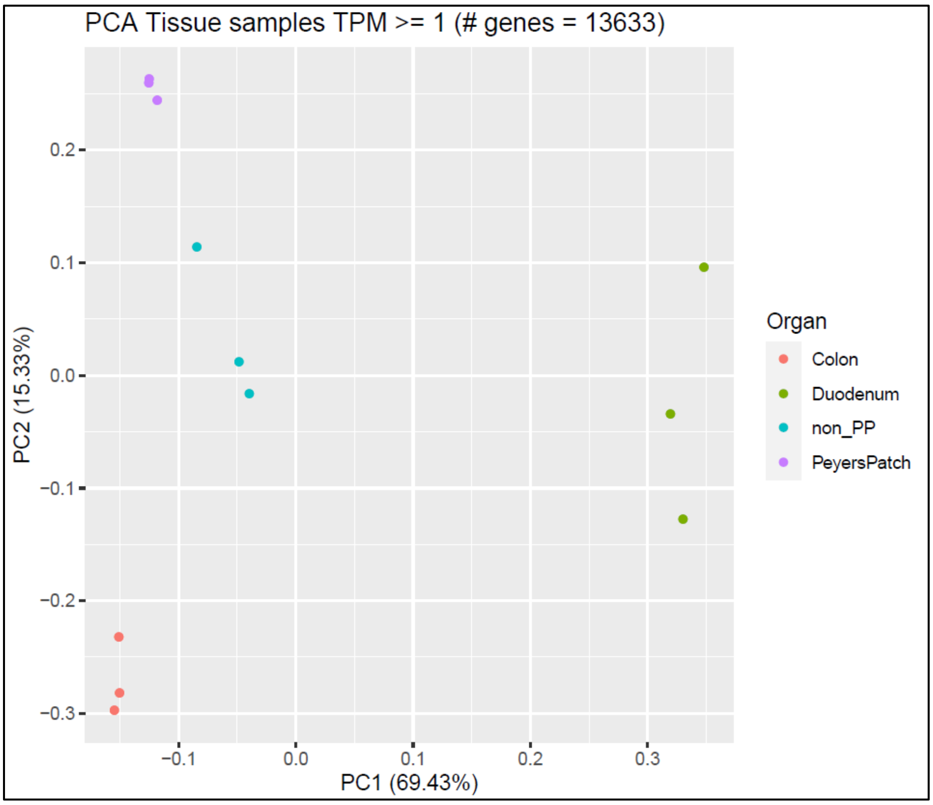
PCA Tissue samples TPM >= 1 (# genes = 13633)
0.2
0.1
0.0
−0.1
−0.2
−0.3
−0.1
0.0
0.1
0.2
0.3
PC2 (15.33%)
PC1 (69.43%)
Organ
Colon
Duodenum
non_PP
PeyersPatch

**C**

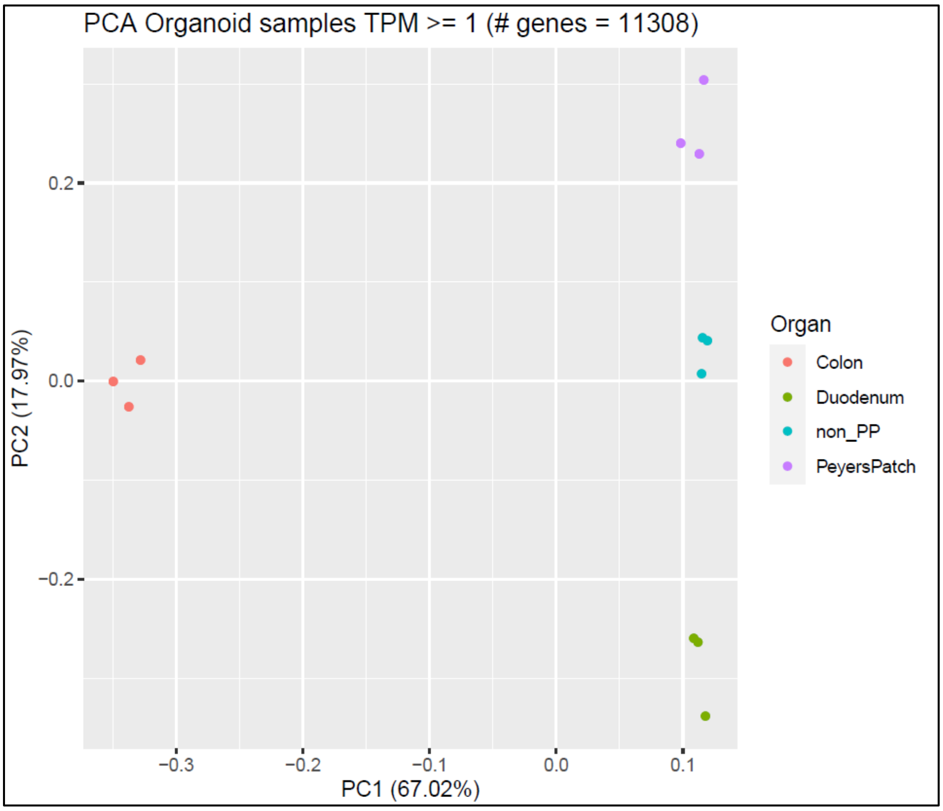

**Supplementary Figure S3**
Transcriptome profiling and clustering of the intestine segments and derived organoids. A. Organoids cluster: colon is grouped more distant from other organoids; intestine segments cluster: duodenum is grouped more distant from other tissues. B. Intestine segments clustering: duodenum is grouped more distant from other tissues, ileum with and without Peyer's Patches (PP) are separated; C: organoids clustering: colon is grouped more distant from other intestine segments, ileum with and without PP are separated

**A**

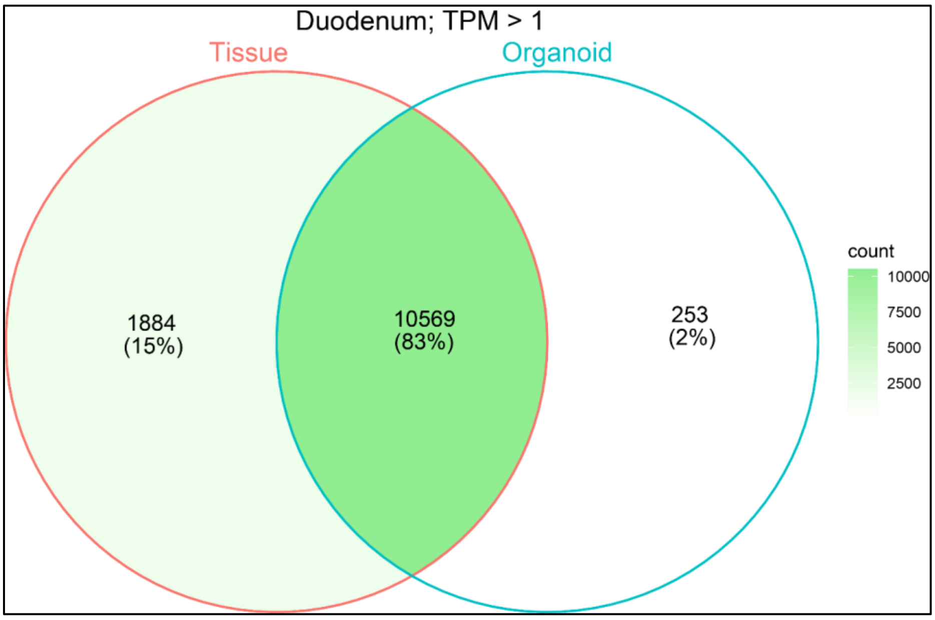

B

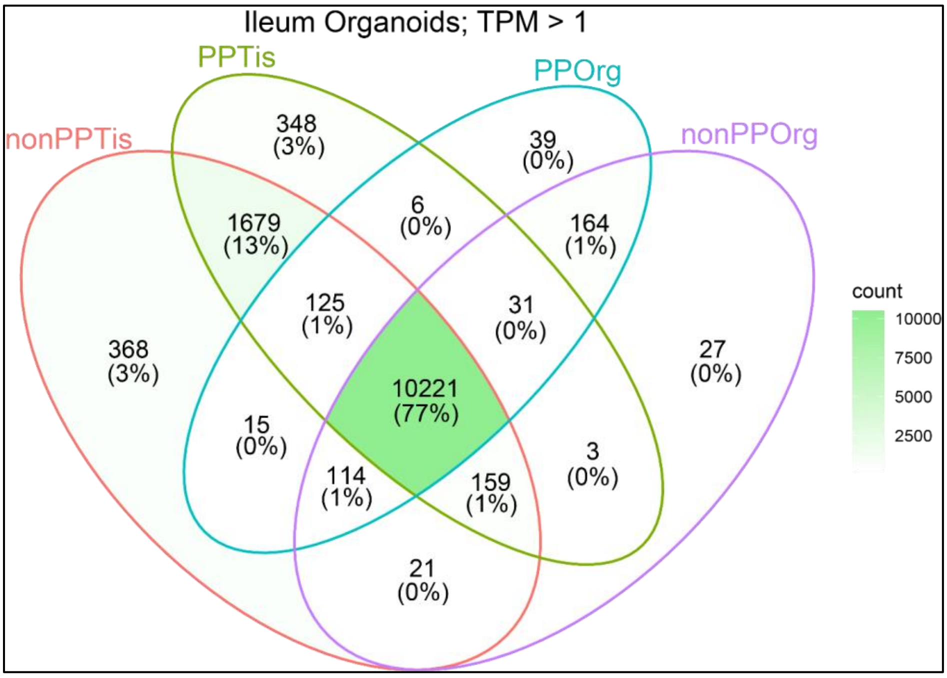
Ileum Organoids; TPM > 1
PPTis
PPOrg
nonPPTis
nonPPOrg
348
(3%)
39
(0%)
6
(0%)
1679
(13%)
164
(1%)
125
(1%)
31
(0%)
368
(3%)
10221
(77%)
27
(0%)
15
(0%)
114
(1%)
159
(1%)
3
(0%)
21
(0%)
count
10000
7500
5000
2500

C

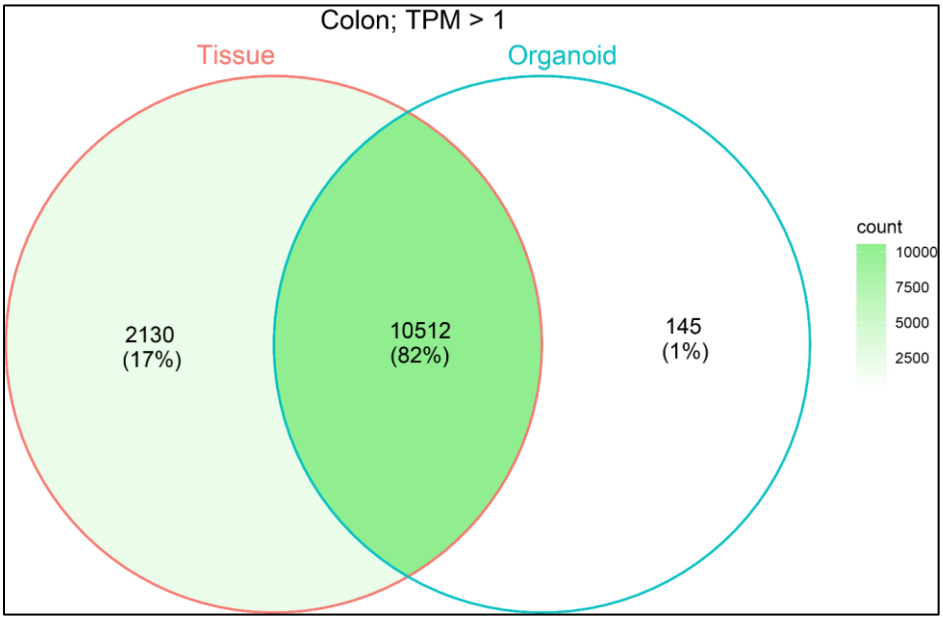
Colon; TPM > 1
Tissue
Organoid
2130
(17%)
10512
(82%)
145
(1%)
count
10000
7500
5000
2500

D

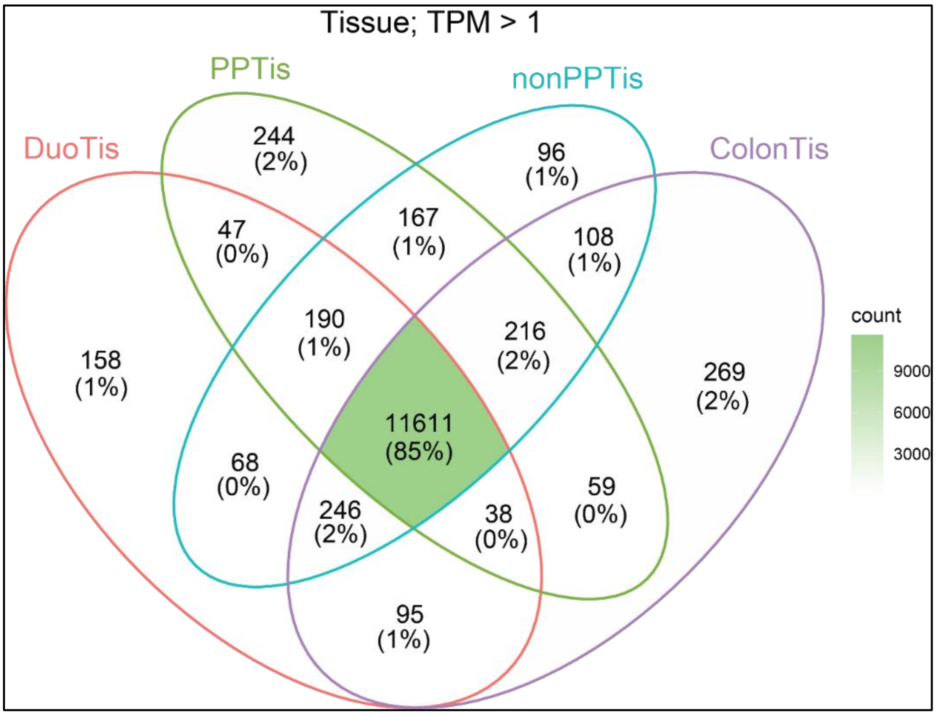

**E**

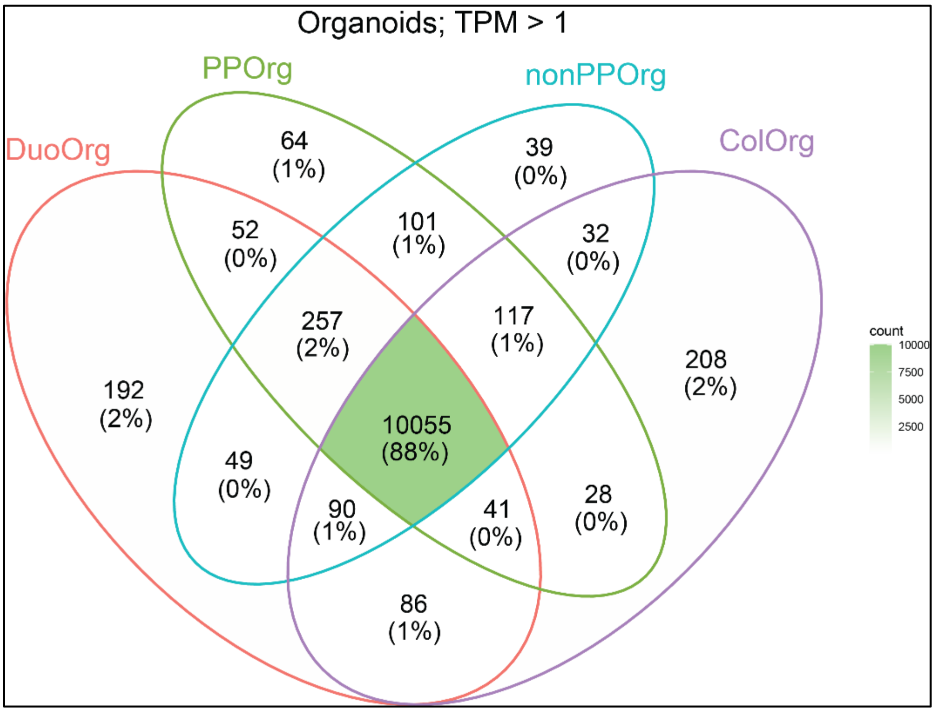

**Supplementary Figure S4**
Organoids and *in vivo* tissue transcriptome profiles share large parts of the transcriptome profiles. A: duodenum; B: ileum; C: colon; D: all intestine segments; E: all organoids.

**Supplementary Table S1**
RNAseq data of the *in vivo* intestine segments and derived organoids. For each intestine segment and each organoid site three independent replicates were sequenced.

| Type[1] | Intestine segment[2] | Replicate | Raw reads | Trimmed reads | Uniquely mapped reads | % Uniquely mapped reads |
|---|---|---|---|---|---|---|
| Org | Colon | 1 | 31,714,381 | 31,533,037 | 30,325,264 | 96.17 |
| Org | Colon | 2 | 32,152,612 | 31,968,860 | 30,691,600 | 96.00 |
| Org | Colon | 3 | 32,206,851 | 32,023,915 | 30,719,448 | 95.93 |
| Org | Duodenum | 2 | 32,512,457 | 32,322,568 | 31,213,528 | 96.57 |
| Org | Duodenum | 3 | 31,554,034 | 31,374,898 | 30,116,407 | 95.99 |
| Org | Duodenum | 4 | 31,392,475 | 31,195,789 | 29,914,739 | 95.89 |
| Org | non_PP | 1 | 32,582,814 | 32,372,380 | 31,220,468 | 96.44 |
| Org | non_PP | 3 | 31,669,021 | 31,477,369 | 30,465,928 | 96.79 |
| Org | non_PP | 4 | 33,314,807 | 33,114,369 | 32,085,627 | 96.89 |
| Org | PP | 2 | 32,233,661 | 32,043,423 | 31,031,362 | 96.84 |
| Org | PP | 3 | 32,044,717 | 31,855,144 | 30,744,931 | 96.51 |
| Org | PP | 4 | 32,059,903 | 31,875,090 | 30,809,448 | 96.66 |
| Tis | Colon | 1 | 32,642,040 | 32,369,185 | 30,974,094 | 95.69 |
| Tis | Colon | 2 | 32,019,705 | 31,750,150 | 30,374,240 | 95.67 |
| Tis | Colon | 3 | 31,904,456 | 31,649,360 | 30,313,945 | 95.78 |
| Tis | Duodenum | 2 | 33,081,572 | 32,825,365 | 31,185,179 | 95.00 |
| Tis | Duodenum | 3 | 32,358,845 | 32,111,419 | 30,504,276 | 95.00 |
| Tis | Duodenum | 4 | 33,339,382 | 33,062,548 | 31,363,681 | 94.86 |
| Tis | non_PP | 1 | 31,817,314 | 31,522,870 | 30,130,298 | 95.58 |
| Tis | non_PP | 3 | 31,286,185 | 30,990,652 | 29,618,055 | 95.57 |
| Tis | non_PP | 4 | 33,403,714 | 33,103,480 | 31,757,602 | 95.93 |
| Tis | PP | 2 | 33,095,632 | 32,778,660 | 31,451,696 | 95.95 |
| Tis | PP | 3 | 31,427,506 | 31,137,830 | 29,880,499 | 95.96 |
| Tis | PP | 4 | 32,407,774 | 32,114,062 | 30,792,923 | 95.89 |

1: Org = organoids, Tis = Tissue (*in vivo*); 2: PP = Peyer's Patches from the ileum intestine segment

**General overview of the RNAseq data:**
Mean ± SD for all samples (organoid and tissue together)

| | |
|---|---|
| Raw input reads | 32,259,200 ± 623,344 |
| Removed basepairs <cutoff | 235,393 ± 47,912 |
| Number of input reads | 32,023,900 ± 612,736 |
| Uniquely mapped reads number | 30,736,900 ± 598,679 |
| Uniquely mapped reads % | 95.98 ± 0.55 |
| Total number of genes | 31,908 |
| Number of expressed genes (TPM > 0) | 15,034 |
| Number of expressed genes (TPM >= 1) | 13,113 |

**Supplementary Table S2**

Genes expected to be specific/indicative for an intestinal location of the expression and RNAseq expression information. Org = organoid; Tiss = tussue; PP = Peyers Patches; Duod = duodenum.

| Gene name | location | Mean TPM_Org Colon | Mean TPM_Org Duod | Mean TPM_ Org non_PP | meanTPM_ OrgPP | Mean TPM_Tiss Colon | Mean TPM_Tiss Duod | Mean TPM_Tissn on_PP | Mean TPM_Ti ssPP |
|---|---|---|---|---|---|---|---|---|---|
| SLC39A 4 | duodenum | 115.490 | 221.030 | 164.320 | 232.553 | 102.657 | 317.000 | 156.197 | 52.693 |
| SLC5A4 | duodenum | 0.520 | 13.790 | 1.693 | 2.933 | 1.017 | 59.703 | 9.887 | 6.867 |
| SLC25A 1 | duodenum | 158.727 | 172.743 | 128.313 | 130.463 | 81.523 | 149.620 | 114.037 | 58.367 |
| SLC37A 4 | duodenum | 28.960 | 33.247 | 24.067 | 27.377 | 37.427 | 63.407 | 53.577 | 22.113 |
| SLC25A 4 | duodenum | 39.467 | 66.663 | 42.137 | 39.003 | 36.383 | 115.570 | 148.430 | 66.730 |
| SLC13A 2 | duodenum | 2.353 | 35.473 | 2.977 | 3.310 | 5.797 | 227.560 | 78.463 | 28.523 |
| LECT2 | duodenum | 0.000 | 0.030 | 0.000 | 0.000 | 0.000 | 7.227 | 0.217 | 0.000 |
| SLC44A 4 | colon | 328.950 | 288.947 | 216.750 | 338.600 | 586.253 | 397.187 | 371.500 | 149.057 |
| GPA33 | colon | 330.573 | 220.937 | 214.847 | 244.400 | 259.747 | 190.740 | 177.870 | 54.990 |
| NR1I3 | colon | 0.290 | 5.010 | 0.423 | 0.353 | 0.550 | 38.947 | 8.490 | 2.580 |
| NOX1 | colon | 0.570 | 0.013 | 0.067 | 0.127 | 26.797 | 0.077 | 0.893 | 1.040 |
| LYZ | jejunum_i leum | 0.000 | 0.000 | 0.000 | 0.000 | 0.000 | 0.000 | 0.000 | 0.000 |
| SLC5A1 | jejunum_i leum | 23.980 | 133.990 | 46.700 | 188.920 | 42.443 | 300.327 | 211.827 | 158.843 |
| DPP4 | jejunum_i leum | 8.883 | 64.390 | 40.907 | 52.463 | 37.943 | 267.747 | 287.043 | 114.973 |
| VIL1 | jejunum_i leum | 323.867 | 575.247 | 745.953 | 1186.530 | 204.247 | 482.343 | 715.867 | 259.457 |
| FABP2 | jejunum_i leum | 0.000 | 116.577 | 6.240 | 12.540 | 1.033 | 336.780 | 427.233 | 202.170 |
| REG4 | jejunum_i leum | 960.363 | 884.163 | 6555.17 7 | 7714.497 | 597.313 | 300.020 | 134.550 | 61.910 |
| KLK1 | Ileum PP | 0.000 | 0.000 | 0.000 | 0.000 | 0.217 | 0.050 | 1.243 | 4.400 |
| BAIAP2 L2 | colon | 81.490 | 92.210 | 172.243 | 230.590 | 99.963 | 93.647 | 279.897 | 85.570 |
| MYO1A | colon | 72.457 | 188.933 | 187.533 | 342.513 | 151.053 | 305.107 | 293.673 | 145.520 |
| PKP2 | colon | 99.503 | 111.910 | 109.110 | 113.047 | 43.520 | 25.830 | 20.787 | 16.247 |
| A2ML1 | colon | 0.577 | 0.073 | 0.293 | 0.107 | 1.840 | 0.083 | 0.047 | 0.023 |
| APOBE C1 | colon | 24.753 | 95.283 | 82.477 | 145.750 | 19.760 | 107.143 | 16.123 | 16.377 |
| PLEKH G6 | colon | 37.297 | 31.383 | 37.203 | 71.673 | 58.260 | 34.667 | 69.030 | 17.913 |
| GMDS | colon | 320.187 | 186.090 | 268.413 | 336.860 | 381.467 | 162.253 | 127.450 | 52.770 |
| TRIM15 | colon | 18.607 | 18.400 | 22.953 | 35.683 | 57.200 | 38.063 | 73.243 | 24.293 |
| GPX2 | colon | 1798.243 | 157.813 | 1178.21 3 | 805.867 | 3367.510 | 45.083 | 159.977 | 89.167 |
| PSEN1 | colon | 206.077 | 110.607 | 137.027 | 186.347 | 117.947 | 76.623 | 97.670 | 58.573 |
| FOXA3 | colon | 4.717 | 8.320 | 13.603 | 20.493 | 18.477 | 13.773 | 4.383 | 2.173 |
| TSPAN1 | colon | 2140.690 | 710.513 | 1010.24 3 | 1708.037 | 3195.127 | 146.100 | 78.053 | 84.027 |
| GCNT3 | colon | 64.237 | 289.293 | 226.333 | 434.043 | 59.647 | 158.740 | 30.273 | 31.177 |
| DUOX2 | colon | 1.907 | 83.623 | 53.320 | 105.613 | 133.653 | 4.310 | 1.797 | 4.770 |
| BMP4 | colon | 3.547 | 5.710 | 5.223 | 7.890 | 17.960 | 9.210 | 11.530 | 12.037 |
| ANXA1 3 | colon | 38.290 | 811.660 | 531.090 | 729.743 | 50.383 | 105.823 | 87.103 | 42.243 |

| | | | | | | | | | |
|---|---|---|---|---|---|---|---|---|---|
| CDH17 | colon | 1320.657 | 480.237 | 841.063 | 971.133 | 493.357 | 286.717 | 586.303 | 179.167 |
| XKR9 | colon | 0.820 | 2.490 | 3.570 | 5.777 | 1.300 | 10.040 | 34.697 | 9.043 |
| GPA33 | colon | 330.573 | 220.937 | 214.847 | 244.400 | 259.747 | 190.740 | 177.870 | 54.990 |
| SLC44A3 | colon | 15.190 | 9.917 | 8.323 | 7.953 | 18.927 | 12.110 | 10.927 | 3.657 |
| CLCA4 | colon | 6.447 | 158.527 | 144.490 | 335.870 | 55.327 | 631.840 | 1138.910 | 324.143 |
| CLCA1 | colon | 294.583 | 60.530 | 835.050 | 1718.303 | 810.620 | 167.460 | 182.227 | 101.880 |
| R3HDML | colon | 0.027 | 0.013 | 0.027 | 0.040 | 0.047 | 0.060 | 0.050 | 0.067 |
| HNF4A | colon | 309.463 | 225.723 | 209.853 | 265.873 | 163.030 | 78.313 | 104.493 | 36.833 |
| MLXIPL | colon | 2.993 | 5.603 | 3.557 | 5.003 | 2.200 | 14.483 | 18.270 | 4.240 |
| TMC5 | colon | 13.140 | 101.003 | 46.243 | 91.010 | 34.563 | 136.707 | 103.843 | 40.787 |
| PROM2 | colon | 13.710 | 267.893 | 135.627 | 155.027 | 0.273 | 0.307 | 0.090 | 0.000 |
| EDAR | colon | 1.037 | 1.017 | 0.343 | 0.753 | 3.017 | 1.463 | 1.440 | 0.867 |
| SLC9A2 | colon | 47.053 | 88.123 | 60.903 | 92.223 | 43.267 | 94.633 | 32.713 | 16.023 |
| ARHGAP25 | colon | 0.017 | 0.020 | 0.020 | 0.030 | 5.890 | 8.417 | 18.993 | 76.990 |
| CORIN | colon | 0.060 | 0.000 | 0.000 | 0.000 | 3.073 | 0.027 | 0.093 | 0.043 |
| CDX2 | colon | 153.360 | 137.997 | 355.070 | 415.243 | 216.083 | 154.633 | 326.080 | 108.943 |
| FREM2 | colon | 0.067 | 0.120 | 0.080 | 0.077 | 0.340 | 0.400 | 0.273 | 0.280 |
| OLFM4 | colon | 8.420 | 147.513 | 23.110 | 10.530 | 2515.853 | 814.247 | 505.817 | 113.863 |
| SLC5A1 | colon | 23.980 | 133.990 | 46.700 | 188.920 | 42.443 | 300.327 | 211.827 | 158.843 |
| HKDC1 | colon | 2.017 | 91.873 | 37.503 | 49.013 | 2.610 | 75.287 | 59.540 | 21.240 |
| KCNH8 | colon | 0.000 | 0.000 | 0.000 | 0.000 | 0.167 | 0.203 | 0.097 | 0.110 |
| LIPH | colon | 93.350 | 124.863 | 101.640 | 144.943 | 31.873 | 34.193 | 39.840 | 15.570 |
| SENP5 | colon | 11.233 | 5.040 | 4.477 | 4.420 | 12.527 | 7.707 | 11.210 | 20.553 |
| MUC13 | colon | 46.210 | 188.770 | 177.240 | 298.073 | 50.397 | 364.110 | 425.737 | 166.053 |
| FANCB | colon | 0.940 | 1.103 | 1.727 | 0.567 | 1.173 | 0.807 | 1.493 | 5.313 |
| HEPH | colon | 99.723 | 191.487 | 221.037 | 259.577 | 56.017 | 96.587 | 78.617 | 29.560 |
| POF1B | colon | 26.927 | 67.557 | 73.843 | 109.713 | 36.690 | 29.673 | 51.500 | 21.663 |
| SLC22A18 | colon | 175.563 | 196.820 | 178.167 | 271.463 | 60.950 | 128.383 | 43.447 | 12.563 |
| LRP4 | colon | 0.070 | 4.880 | 0.457 | 1.120 | 10.950 | 59.190 | 13.800 | 4.930 |
| USH1C | colon | 76.007 | 77.637 | 106.037 | 130.313 | 108.430 | 148.353 | 170.143 | 51.707 |
| B3GNT3 | colon | 231.917 | 47.347 | 90.563 | 94.613 | 129.893 | 8.087 | 4.957 | 8.370 |
| SLC12A2 | colon | 38.043 | 34.190 | 100.700 | 57.330 | 119.460 | 67.627 | 50.303 | 23.890 |
| CDX1 | colon | 53.423 | 47.133 | 62.963 | 56.377 | 34.710 | 7.327 | 10.290 | 3.643 |
| TMPRSS4 | colon | 136.257 | 14.040 | 26.313 | 44.393 | 79.257 | 1.863 | 1.760 | 0.793 |
| TMEM45B | colon | 223.533 | 293.283 | 371.513 | 450.020 | 130.907 | 216.653 | 300.193 | 75.683 |
| ST14 | colon | 516.697 | 262.023 | 369.193 | 508.740 | 331.277 | 133.770 | 199.327 | 77.733 |
| PIGR | colon | 23.440 | 48.200 | 0.877 | 2.147 | 148.477 | 833.793 | 21.463 | 20.407 |
| MYO7B | colon | 156.500 | 120.313 | 106.360 | 138.160 | 201.003 | 122.420 | 183.023 | 65.070 |
| VIL1 | colon | 323.867 | 575.247 | 745.953 | 1186.530 | 204.247 | 482.343 | 715.867 | 259.457 |
| IHH | colon | 182.943 | 32.083 | 32.343 | 76.443 | 93.817 | 32.550 | 48.373 | 21.717 |
| CFTR | colon | 20.790 | 14.220 | 18.323 | 16.403 | 34.720 | 64.193 | 60.620 | 19.540 |
| EVX1 | colon | 1.180 | 0.000 | 0.000 | 0.000 | 1.197 | 0.000 | 0.000 | 0.000 |
| AXIN2 | colon | 5.223 | 3.073 | 5.843 | 5.917 | 12.267 | 12.480 | 28.327 | 7.950 |
| KRT20 | colon | 964.810 | 1761.797 | 2766.043 | 3578.103 | 461.373 | 1000.170 | 404.500 | 263.613 |

| | | | | | | | | | |
|---|---|---|---|---|---|---|---|---|---|
| HOXB8 | colon | 9.953 | 0.000 | 16.387 | 13.207 | 2.097 | 0.187 | 5.493 | 3.297 |
| HOXB6 | colon | 57.330 | 0.117 | 46.440 | 34.763 | 140.910 | 1.700 | 172.287 | 56.507 |
| ADAMTS13 | colon | 0.070 | 0.143 | 0.007 | 0.063 | 0.837 | 1.997 | 1.197 | 1.797 |
| DAPK2 | colon | 0.583 | 1.080 | 0.153 | 0.293 | 1.767 | 4.997 | 3.307 | 2.133 |
| SATB2 | colon | 85.807 | 0.130 | 13.743 | 16.447 | 70.513 | 0.097 | 20.487 | 7.973 |
| SLC26A3 | colon | 447.947 | 89.780 | 73.573 | 169.437 | 200.433 | 8.173 | 157.630 | 56.017 |
| DPEP1 | colon | 6.027 | 3.050 | 18.280 | 53.693 | 196.507 | 43.280 | 1827.820 | 426.897 |
| LGALS4 | colon | 173.623 | 160.967 | 187.433 | 228.323 | 172.787 | 118.887 | 76.540 | 34.667 |
| TMPRSS3 | colon | 0.063 | 0.293 | 0.080 | 0.043 | 0.870 | 0.740 | 0.043 | 0.020 |
| RAB20 | colon | 8.113 | 8.217 | 3.453 | 4.377 | 13.313 | 14.740 | 39.943 | 14.250 |
| TFF3 | colon | 335.200 | 533.410 | 2077.503 | 4903.343 | 1057.287 | 1452.040 | 609.570 | 268.633 |
| GRM8 | colon | 0.183 | 0.000 | 0.000 | 0.000 | 0.277 | 0.043 | 0.027 | 0.197 |
| IL1RL2 | colon | 0.000 | 0.000 | 0.000 | 0.000 | 0.427 | 0.030 | 0.043 | 0.040 |
| MT1A | colon | 2.967 | 68.383 | 11.710 | 32.963 | 92.497 | 214.033 | 86.427 | 39.140 |
| SGK2 | colon | 32.443 | 31.127 | 5.760 | 8.297 | 76.700 | 86.800 | 8.723 | 2.133 |
| PTPRH | colon | 56.807 | 117.007 | 99.277 | 172.160 | 80.107 | 128.033 | 97.443 | 45.800 |
| ETV3 | colon | 27.427 | 37.127 | 34.453 | 56.070 | 24.480 | 33.143 | 25.347 | 23.740 |
| TMEM54 | colon | 635.543 | 319.390 | 448.117 | 573.540 | 879.243 | 210.750 | 126.580 | 73.153 |
| EPHB3 | colon | 4.100 | 3.060 | 4.370 | 4.783 | 9.160 | 8.597 | 4.493 | 7.647 |
| SIM2 | colon | 0.067 | 0.000 | 0.000 | 0.000 | 0.037 | 0.013 | 0.013 | 0.027 |
| MYH14 | colon | 234.600 | 194.997 | 164.977 | 237.867 | 398.303 | 149.483 | 195.007 | 67.587 |
| MST1R | colon | 244.363 | 90.663 | 100.587 | 114.010 | 91.893 | 45.727 | 42.347 | 16.277 |
| POU2F3 | colon | 0.543 | 0.123 | 0.437 | 0.927 | 5.177 | 1.260 | 6.057 | 14.067 |
| SLC6A20 | colon | 15.367 | 25.047 | 19.320 | 30.953 | 33.133 | 10.463 | 23.480 | 15.017 |
| DNMT3A | colon | 7.467 | 7.330 | 7.713 | 7.807 | 14.880 | 13.250 | 17.357 | 10.293 |
| REN | colon | 0.000 | 0.000 | 0.000 | 0.000 | 0.030 | 0.000 | 0.030 | 0.030 |
| AQP8 | colon | 3.840 | 3.697 | 2.030 | 11.957 | 19.733 | 71.207 | 29.757 | 13.640 |
| SLC17A4 | colon | 0.000 | 0.000 | 0.000 | 0.000 | 0.000 | 0.000 | 0.000 | 0.000 |
| RSPO1 | colon | 0.000 | 0.380 | 3.117 | 4.603 | 0.440 | 0.460 | 0.840 | 0.337 |
| CYP2S1 | colon | 11.320 | 137.610 | 7.510 | 10.627 | 9.710 | 11.610 | 5.713 | 3.460 |
| ASCL2 | colon | 0.350 | 2.463 | 1.420 | 0.427 | 26.417 | 10.450 | 7.973 | 3.367 |
| COL8A2 | colon | 0.027 | 0.020 | 0.013 | 0.020 | 1.540 | 0.177 | 0.620 | 0.163 |
| RNF186 | colon | 0.970 | 1.430 | 1.813 | 2.907 | 3.980 | 15.860 | 2.703 | 1.347 |
| CLDN3 | colon | 267.270 | 134.713 | 203.980 | 453.757 | 423.353 | 149.483 | 77.343 | 58.010 |
| OVOL2 | colon | 2.680 | 4.847 | 2.113 | 2.787 | 7.947 | 5.943 | 5.773 | 1.487 |
| TMEM92 | colon | 39.177 | 33.837 | 45.267 | 73.047 | 32.633 | 36.517 | 59.653 | 21.373 |
| CLDN2 | colon | 20.600 | 115.657 | 109.057 | 190.540 | 68.973 | 374.403 | 250.560 | 67.500 |
| TRIM7 | colon | 3.060 | 5.830 | 4.050 | 4.317 | 2.430 | 2.593 | 3.730 | 1.423 |
| IL22RA1 | colon | 26.663 | 32.013 | 25.007 | 38.023 | 23.360 | 44.537 | 88.057 | 23.153 |
| HOXB9 | colon | 22.570 | 0.013 | 20.780 | 21.110 | 31.843 | 0.223 | 14.637 | 5.310 |
| KCNQ1 | colon | 270.280 | 249.360 | 204.890 | 254.207 | 154.807 | 146.417 | 30.990 | 12.487 |
| GPR35 | colon | 33.597 | 24.077 | 18.500 | 26.633 | 30.080 | 23.297 | 20.290 | 15.567 |
| REG4 | colon | 960.363 | 884.163 | 6555.177 | 7714.497 | 597.313 | 300.020 | 134.550 | 61.910 |

| | | | | | | | | | |
|---|---|---|---|---|---|---|---|---|---|
| NELL2 | ileum | 0.010 | 0.010 | 0.010 | 0.000 | 1.167 | 1.360 | 1.603 | 3.173 |
| FURIN | ileum | 68.700 | 60.087 | 55.263 | 74.187 | 87.457 | 104.980 | 73.440 | 50.853 |
| C15orf39 | ileum | 2.003 | 4.073 | 6.243 | 6.983 | 4.190 | 6.157 | 3.937 | 5.293 |
| IPO4 | ileum | 42.847 | 40.760 | 45.287 | 40.287 | 46.123 | 33.890 | 34.627 | 30.753 |
| CLPTM1 | ileum | 72.703 | 96.977 | 88.570 | 103.843 | 67.417 | 110.113 | 128.907 | 65.010 |
| MEP1B | ileum | 1.330 | 7.140 | 15.123 | 33.580 | 11.063 | 144.747 | 981.777 | 286.040 |
| EPHA7 | ileum | 0.010 | 0.000 | 0.000 | 0.003 | 1.460 | 1.177 | 2.277 | 1.537 |
| TPBG | ileum | 6.927 | 2.997 | 2.300 | 2.473 | 3.237 | 1.677 | 2.477 | 6.687 |
| BMP4 | ileum | 3.547 | 5.710 | 5.223 | 7.890 | 17.960 | 9.210 | 11.530 | 12.037 |
| OSR2 | ileum | 0.097 | 0.287 | 59.093 | 86.953 | 0.143 | 0.920 | 33.873 | 16.067 |
| CDH17 | ileum | 1320.657 | 480.237 | 841.063 | 971.133 | 493.357 | 286.717 | 586.303 | 179.167 |
| SLC39A1 | ileum | 40.377 | 36.527 | 36.290 | 43.143 | 58.200 | 49.747 | 48.310 | 35.703 |
| SYPL2 | ileum | 0.773 | 0.423 | 0.540 | 0.367 | 0.477 | 0.957 | 0.690 | 1.080 |
| CLCA1 | ileum | 294.583 | 60.530 | 835.050 | 1718.303 | 810.620 | 167.460 | 182.227 | 101.880 |
| CLCN7 | ileum | 12.377 | 10.747 | 10.767 | 13.270 | 23.777 | 20.317 | 52.497 | 27.337 |
| ABCA3 | ileum | 15.130 | 5.263 | 9.090 | 14.093 | 16.843 | 5.470 | 12.140 | 22.890 |
| ZNF512 | ileum | 1.263 | 0.937 | 0.700 | 0.637 | 6.957 | 7.413 | 9.987 | 15.283 |
| APOB | ileum | 0.900 | 50.757 | 20.733 | 48.513 | 0.520 | 348.123 | 440.283 | 75.923 |
| SLIT2 | ileum | 0.010 | 0.033 | 0.033 | 0.027 | 0.987 | 0.497 | 0.777 | 0.440 |
| EDNRA | ileum | 0.000 | 0.007 | 0.007 | 0.000 | 3.347 | 2.470 | 2.677 | 0.877 |
| MTTP | ileum | 0.027 | 159.013 | 12.770 | 31.353 | 0.587 | 373.023 | 320.710 | 78.457 |
| MMRN1 | ileum | 0.000 | 0.000 | 0.007 | 0.000 | 8.547 | 5.913 | 9.757 | 3.757 |
| SNCA | ileum | 0.000 | 0.000 | 0.000 | 0.073 | 1.007 | 1.987 | 1.567 | 1.183 |
| TNFRSF19 | ileum | 0.123 | 0.153 | 0.400 | 0.363 | 0.647 | 1.680 | 0.883 | 1.527 |
| SPRY2 | ileum | 87.007 | 35.630 | 18.843 | 31.873 | 50.247 | 23.643 | 34.297 | 9.520 |
| DKK1 | ileum | 0.057 | 0.000 | 0.070 | 0.020 | 0.107 | 0.097 | 0.363 | 2.100 |
| SCD | ileum | 887.490 | 540.923 | 335.257 | 307.507 | 75.843 | 6.897 | 8.303 | 29.617 |
| CNNM2 | ileum | 5.267 | 0.420 | 0.237 | 0.490 | 26.517 | 0.787 | 1.663 | 1.287 |
| SEC61A1 | ileum | 127.553 | 104.430 | 123.280 | 132.727 | 126.680 | 174.157 | 113.297 | 106.583 |
| SI | ileum | 0.053 | 0.327 | 26.190 | 73.997 | 0.703 | 3.120 | 282.537 | 151.080 |
| TFRC | ileum | 184.447 | 155.010 | 51.210 | 41.420 | 72.660 | 129.210 | 22.453 | 39.880 |
| PAX6 | ileum | 0.213 | 0.153 | 0.530 | 0.940 | 0.673 | 0.507 | 1.250 | 2.677 |
| CALR | ileum | 944.383 | 729.133 | 900.720 | 923.003 | 884.257 | 1151.980 | 1330.343 | 869.837 |
| ENC1 | ileum | 8.243 | 3.693 | 5.067 | 7.437 | 3.493 | 7.497 | 2.773 | 1.593 |
| TCOF1 | ileum | 9.893 | 16.843 | 15.580 | 8.200 | 20.197 | 18.693 | 25.773 | 46.120 |
| SPCS2 | ileum | 130.573 | 104.277 | 116.533 | 114.320 | 145.677 | 147.727 | 141.400 | 155.090 |
| GCG | ileum | 41.390 | 1.673 | 242.943 | 520.667 | 153.673 | 0.430 | 351.683 | 85.443 |
| ITGAV | ileum | 20.427 | 56.890 | 29.307 | 43.340 | 31.097 | 32.067 | 29.733 | 20.547 |
| COL5A2 | ileum | 0.393 | 0.363 | 0.517 | 0.400 | 57.240 | 33.280 | 52.280 | 46.527 |
| KIF1A | ileum | 0.640 | 0.237 | 0.387 | 0.580 | 3.180 | 1.303 | 1.330 | 0.440 |
| FGF10 | ileum | 0.000 | 0.007 | 0.000 | 0.000 | 1.777 | 0.427 | 0.227 | 0.210 |
| GAA | ileum | 39.377 | 16.467 | 11.167 | 14.400 | 20.823 | 14.423 | 12.887 | 16.493 |
| SEC14L1 | ileum | 26.713 | 32.217 | 18.373 | 19.683 | 25.100 | 46.010 | 32.320 | 47.783 |
| DHCR7 | ileum | 59.657 | 108.833 | 58.130 | 48.920 | 69.707 | 55.540 | 31.543 | 24.843 |
| SLC4A2 | ileum | 40.780 | 37.097 | 24.500 | 21.273 | 73.990 | 62.450 | 70.707 | 40.163 |

| | | | | | | | | | |
|---|---|---|---|---|---|---|---|---|---|
| SLC26A3 | ileum | 447.947 | 89.780 | 73.573 | 169.437 | 200.433 | 8.173 | 157.630 | 56.017 |
| PHF6 | ileum | 11.247 | 11.057 | 10.467 | 8.333 | 15.350 | 14.027 | 28.730 | 73.187 |
| CCDC9 | ileum | 43.710 | 47.130 | 47.027 | 58.657 | 54.167 | 47.583 | 66.060 | 83.530 |
| CPT1C | ileum | 0.013 | 0.050 | 0.090 | 0.010 | 1.590 | 0.710 | 1.523 | 4.617 |
| STX12 | ileum | 53.413 | 72.980 | 63.103 | 76.607 | 47.363 | 60.110 | 45.857 | 30.637 |
| ALDH1A1 | ileum | 105.483 | 7426.123 | 1557.893 | 1653.293 | 50.830 | 1957.893 | 216.827 | 128.590 |
| ZNF398 | ileum | 4.473 | 6.700 | 4.960 | 5.613 | 6.847 | 9.947 | 11.007 | 15.450 |
| GANAB | ileum | 168.883 | 121.993 | 129.653 | 114.900 | 139.973 | 110.413 | 114.887 | 117.020 |
| NDST1 | ileum | 9.100 | 9.463 | 8.777 | 7.670 | 24.960 | 47.463 | 40.790 | 27.203 |
| PCDH18 | ileum | 0.063 | 0.007 | 0.007 | 0.013 | 6.057 | 3.590 | 6.123 | 2.157 |
| KREMEN2 | ileum | 1.337 | 1.670 | 1.443 | 1.323 | 0.850 | 0.017 | 0.550 | 2.330 |
| KITLG | ileum | 10.620 | 7.613 | 11.243 | 10.773 | 14.327 | 22.300 | 18.087 | 8.713 |
| SPIN1 | ileum | 24.457 | 28.187 | 32.010 | 34.200 | 32.023 | 25.397 | 33.757 | 33.657 |
| PTDSS2 | ileum | 33.963 | 36.763 | 28.350 | 35.897 | 57.143 | 48.443 | 49.517 | 30.780 |
| DUSP6 | ileum | 108.533 | 66.077 | 90.670 | 137.293 | 33.237 | 41.410 | 42.323 | 42.243 |
| ALDOB | ileum | 5.700 | 420.937 | 35.790 | 98.380 | 65.200 | 1597.533 | 1020.453 | 370.523 |
| PCDH9 | ileum | 0.000 | 0.000 | 0.000 | 0.000 | 0.223 | 0.090 | 0.147 | 0.117 |
| M6PR | ileum | 64.477 | 75.143 | 62.767 | 62.190 | 65.407 | 61.630 | 119.340 | 87.850 |
